# Reproducibility metrics for CRISPR screens

**DOI:** 10.1101/2022.02.19.480892

**Authors:** Maximilian Billmann, Henry N. Ward, Michael Aregger, Michael Costanzo, Brenda J. Andrews, Charles Boone, Jason Moffat, Chad L. Myers

## Abstract

CRISPR screens are used extensively to systematically interrogate the phenotype-to-genotype problem. In contrast to early CRISPR screens, which defined core cell fitness genes, most current efforts now aim to identify *context-specific* phenotypes that differentiate a cell line, genetic background or condition of interest, such as a drug treatment. While CRISPR-related technologies have shown great promise and a fast pace of innovation, a better understanding of standards and methods for quality assessment of CRISPR screen results is crucial to guide technology development and application. Specifically, many commonly used metrics for quantifying screen quality do not accurately measure the reproducibility of context-specific hits. We highlight the importance of reporting reproducibility statistics that directly relate to the purpose of the screen and suggest the use of metrics that are sensitive to context-specific signal.

---

CRISPR screens see widespread use in the functional genomics community for interrogating gene function. Most prominently, loss-of-function CRISPR screens measure how the perturbation of each individual gene, across a library of targeted genes, affects cell fitness within a pool of cells. Each gene’s measurement in essence is composed of three elements: a fitness effect common to all cell types, a fitness effect specific to the biological context of the experiment (including cell type), and measurement error. Focusing on the first element, CRISPR screens completed across hundreds of different human cell types have now definitively identified the core genes that are essential across many cell types (Meyers et al., 2017). Given that the core essential genes have been well-established, many CRISPR loss-of-function screens now focus on identifying genes that are essential in specific contexts, including different cell types, genetic backgrounds, or environmental conditions (Dempster et al., 2019; DeWeirdt et al., 2020; Han et al., 2017; Meyers et al., 2017; Najm et al., 2018; Shen et al., 2017; Tsherniak et al., 2017). Context-specific gene essentiality is important to explore because it can potentially help to guide functional annotation of the majority of genes in the human genome or elucidate disease mechanisms and therapy possibilities. However, the reproducibility of the *context-specific* effects discovered by CRISPR screens is inconsistently reported. Biological replicate screen reproducibility is typically reported using correlation measures on the level of normali ed readcount data or fitness effects. At those processing steps, the data largely reflects covariation due to the gRNA representation in the library and/or the consensus (not context-specific) gene essentiality, respectively – neither provides an accurate estimate of the reproducibility of context-specific effects, which is often the main focus of the screen. Moreover, the interpretation of the commonly used metric, a correlation coefficient, is unclear due to the typical sparsity of effects in such screens.

To illustrate our point, we assess alternative reproducibility metrics across data processing levels of differential genome-wide CRISPR-Cas9 screens to identify genetic interactions (GI) with the fatty acid synthase (FASN) (Aregger et al., 2020). Firstly, we report the Pearson correlation coefficient (PCC) between independently replicated screens at the following points of data processing: starting gRNA abundance, end gRNA abundance, a fitness score reflecting the log-fold-change (LFC) between the end and starting gRNA abundance, the context-specific effect as measured by the differential LFC (dLFC; raw GI score), and finally, a fully normali ed dLFC score (expressed as the qGI score, see Aregger et al. 2020 for details) (Figure 1a).

**Figure 1:**
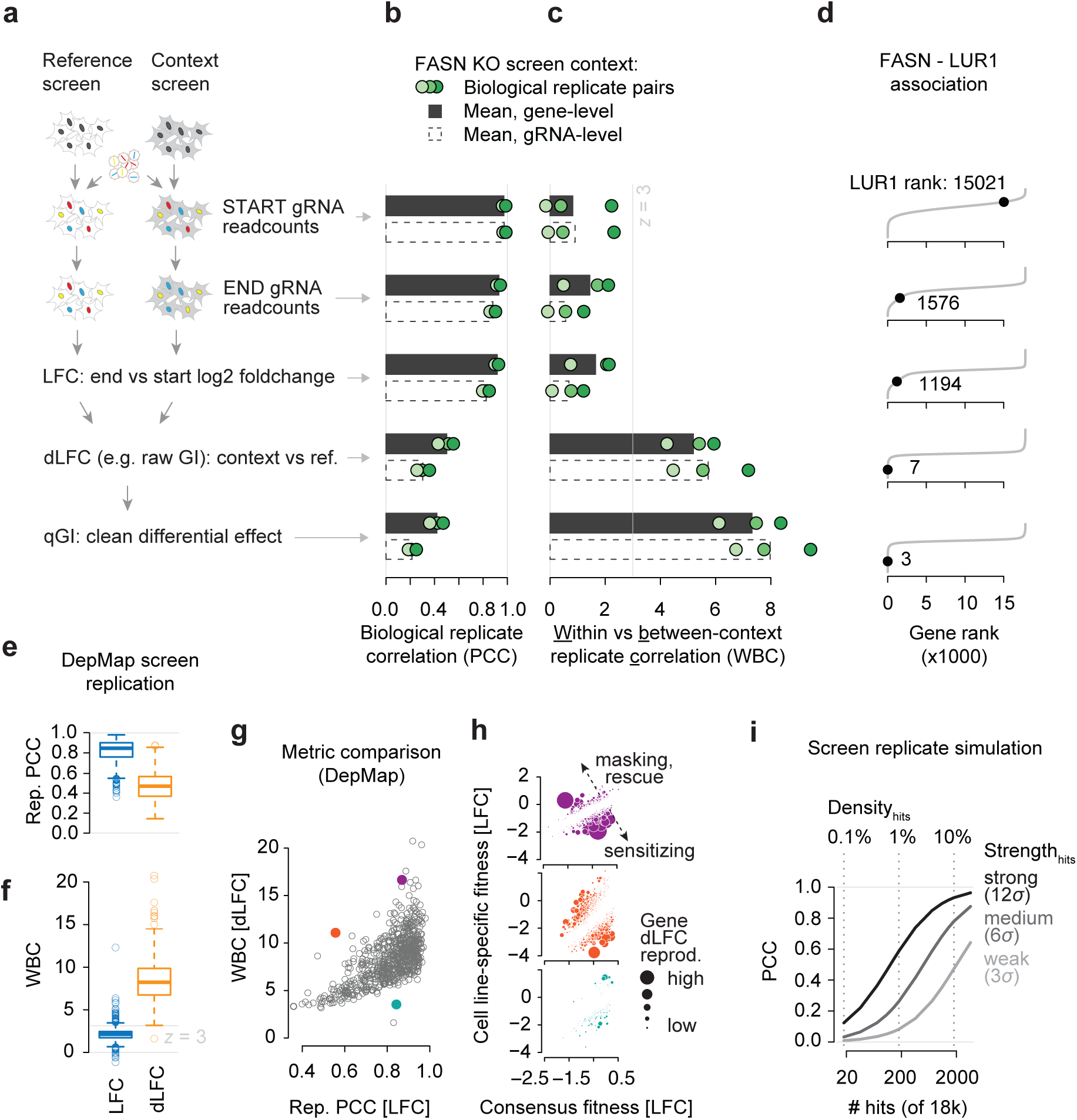
Reproducibility metrics for context-specific signal in CRISPR screens. **(a)** Summary of the context-specific CRISPR-Cas9 screening and differential effect identification process. **(b)** Between-replicate Pearson correlation coefficients (PCC) for start and endpoint readcount data, the log2 foldchange (LFC) thereof and the differential LFC (dLFC) and qGI scores. Bars represent the mean of the three pairwise comparisons, dots represent the individual pairs. Screens were independently performed (starting from preparation and transfection of the gRNA library). **(c)** Within-FASN KO replicate to between FASN KO and non-FASN KO screen ratio of PCCs (WBC; see methods for details). Bars and dots represent the same as explained in (b). **(d)** Ranking of LUR1 (previous C12orf49) among the 17,804 genes screened in FASN KO cells at each data processing step as defined in Aregger et al. 2020. Ranks are means of the three biological replicates. **(e, f)** Between-replicate PCC (e) and WBC (f) of LFC and dLFC data from DepMap genome-wide screens in 693 cell lines. **(g)** Comparison of between screen replicate PCCs for each of the 693 DepMap screens with the WBC. The three cell lines shown in (h) are highlighted. **(h)** Reproducibility of dLFC effects in three cell lines with different sets of replicate PCCs and WBCs. The consensus fitness is the per-gene mean LFC value across all replicates and 693 cell lines. The cell line-specific fitness is the per-gene LFC measured in each given cell line (SKBR3 = violet, HCC1187 = dark orange, MEL202 = cyan). Circle size corresponds to the dLFC product between replicate screens. **(i)** PCC between simulated screening data with normally distributed noise at increasing numbers of hits with weak, medium and strong amplitude. Hit strength is defined as a multiple of the standard deviation of the noise distribution (δ).

Within-context (FASN KO) replicate correlations were highest for starting readcounts (r = 0.97 for gRNA, r = 0.97 for gene-level measures), reflecting that the gRNA library distribution is reproducible (Figure 1b). Removing this library effect from the endpoint readcounts to obtain LFC fitness values also results in high PCCs at the gene-level (r = 0.92) and slightly lower gRNA-level PCCs of 0.82 (Figure 1b). This shows that both unwanted technical features of the experiment and general fitness effects are highly reproducible. However, in this context, we aim to identify GIs with FASN, i.e. FASN-specific fitness effects, and thus, all of the measures above fail to measure the reproducibility associated with the focus of our screen. Once dLFC values are computed between the FASN KO query and wild-type reference screens, the PCC between replicate screens drops substantially to 0.3 (gRNA) and 0.5 (gene) and further to 0.21 (gRNA) and 0.42 (gene) when experimental artifacts are computationally normali ed, which is reflected in the qGI score (Figure 1b). In summary, the replicate correlations decrease with more accurate quantification of the biological signal of interest, which is context-specific effects (in this case, GIs). Importantly, fitness score (LFC)-based replicate correlations cannot approximate context-specific effect reproducibility (Figure S1a-c), and such comparisons are particularly problematic for comparative evaluation of data from different sets of genes or different cell models.

To illustrate why focusing on the appropriate screen statistics is important for reporting reproducibility, we further analy ed the FASN KO, but compared it to five non-FASN KO screens. Under the simple assumption that biological replicates of the same genetic screen should exhibit more similarity than genetic screens with different query mutations, we computed a score that captures the similarity of two replicates of the same screen relative to the similarity of different screens, which we will subsequently refer to as the Within-vs-Between context replicate Correlation (WBC) score (see methods). Despite their high correlations, readcounts and LFC data did not distinguish within-context (same KO) replicates from between-context (different KO) pairs (Figure 1c), confirming that the high similarity does not indicate reproducibility of the main quantity of interest. In contrast, despite low within-context replicate PCCs, the dLFC measure exhibited strong WBC scores (z > 3), and these were further improved in the qGI score (Figure 1c). Moreover, only dLFC and qGI scores identified the recently discovered link between FASN and LUR1 (Figure 1d) (Aregger et al., 2020).

To demonstrate the generality of our findings beyond genetic interaction screens, we performed a similar analysis on the reproducibility of cell line-specific effects in 693 screens within the Cancer Dependency Map(Meyers et al., 2017; Tsherniak et al., 2017). Again, we found that the within-context (same cell line) replicate correlation decreased substantially between the fitness effect (LFC; mean r = 0.81) and the cell line specific deviation from the consensus fitness profile (dLFC; mean r = 0.47) (Figure 1e). In contrast, the WBC score indicates that the dLFC metric reflects context-specific signal with much higher quality (Figure 1f), which is the main goal in building such a cancer dependency map. Specifically, significant (z > 3) and highly significant (z > 5) reproducibility scores for cell line-specific effects are found for 99.9% and 91.3% of all cell lines by using the dLFC metric, respectively, while only 10.1% and 1.2% of cell lines reach the same significance level when using the LFC metric (Figure 1f). A metric like the WBC score provides additional resolution compared to simple correlation measures, emphasi ing the reproducibility of context-specific effects (Figure 1g, h). We note that the antagonistic relationship between replicate correlation and data normali ation extends to non-CRISPR screens, including the most comprehensive GI data to date, recorded in yeast (Figure S2a, b) (Costanzo et al., 2021, 2016).

Perhaps one of the reasons why informative reproducibility measures are not consistently reported is that they tend to be relatively low, which may be viewed as evidence for poor quality data. How high should we expect replicate correlation to be for a high-quality CRISPR screen? Global metrics such as the PCC are highly impacted by the sparsity of the signal one is measuring, and in most cases, one would expect context-specific genetic effects to be rare (Aregger et al., 2020; Costanzo et al., 2016; Tsherniak et al., 2017). For instance, simulated genome-wide screening data (∼18k genes) with normally distributed noise and 18 (0.1% density) or 180 (1% density) strong (12σ) true hits, densities typical of context-specific screens, would result in a PCC of 0.13 and 0.59, respectively. In contrast, a hit-density of 10%, which is more typical for pure fitness phenotypes in genome-wide screens, results in a PCC of 0.93 (Figure 1i). Thus, we should expect low PCC measures in genome-scale screens even where a small number of hits are highly reproducible due to the sparsity of context-specific fitness effects.

In conclusion, we highlight the importance of reporting appropriate reproducibility statistics for CRISPR screens. Biological replicate screens should be performed to establish the quality of data in any screening context, and importantly, the reported statistics should directly relate to the purpose of the screen. In addition to standard correlation measures, we suggest the use of additional metrics, such as the WBC, that are sensitive to context-specific signal.

**Figure S1:**
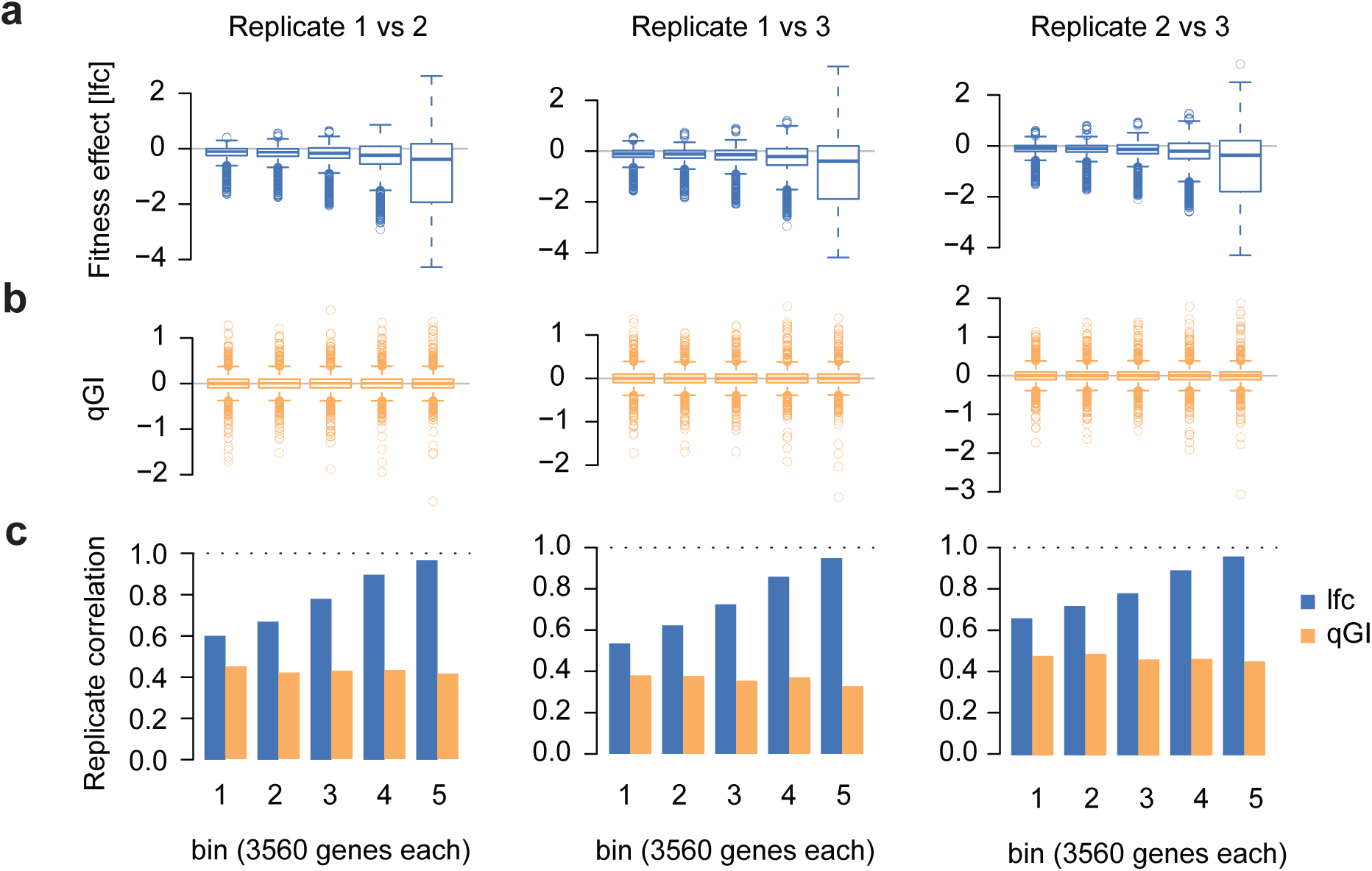
Relation of qGI and lfc-based screen replicate correlation. **(a)** Fitness effects of genes in equally sized bins. The fitness effects represent the mean LFC scores across the two FASN KO replicates screens indicated. **(b)** qGI effects of genes in equally sized bins. qGI scores represent the mean across the two FASN KO screens indicated. **(c)** Replicate Pearson correlation coefficients per bin based on LFC (blue) and qGI (orange) scores. Each bin contains exactly 3560 genes that were all taken from the same screen (see methods).

**Figure S2:**
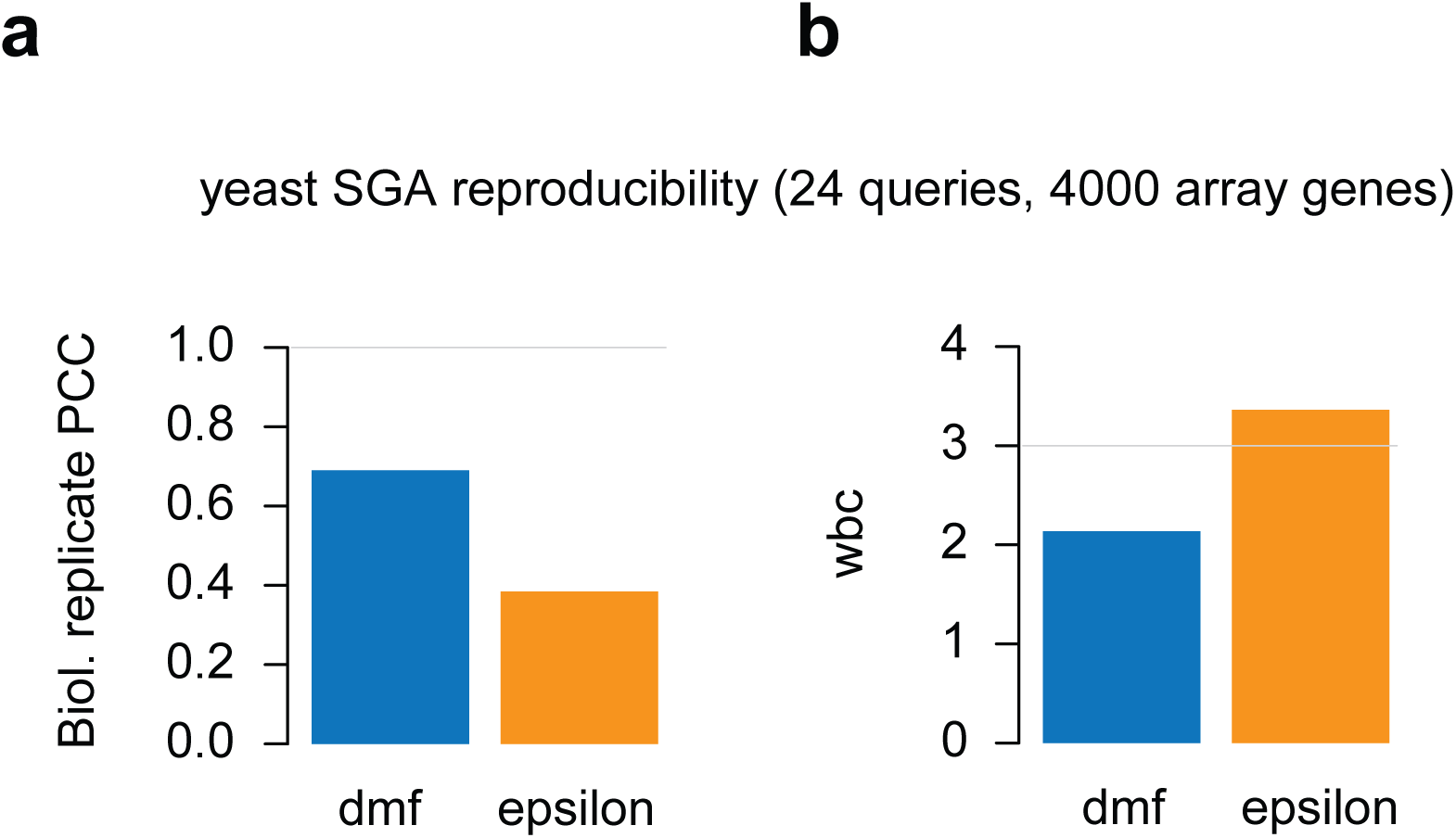
Reproducibility metrics applied to yeast SGA genetic interaction data. **(a)** Between-biological replicate correlation of double mutant fitness(dmf) and genetic interaction scores (epsilon). The mean Pearson correlation coefficient between 24 query genes screened in biological duplicates against ∼4000 array genes is shown. **(b)** WBC scores show the difference between each of the query screens contrasted to the remaining 23 queries by using dmf and epsilon scores.

## Supplementary Information

### Methods

#### Replicate correlation of genome-wide CRISPR-Cas9 screens in HAP1 FASN KO cells

HAP1 genome-wide CRISPR-Cas9 screening data was taken from Aregger et al. 2020. The three biological replicates were independently screened (including gRNA library preparation and transfection). The gRNA-level comparisons use values from 70,006 gRNAs that had an initial abundance at the start of the experiment of at least 40 readcounts. The gene-level comparisons use values from 17,804 genes, considering a gene whenever at least two gRNA sequences pass all QC thresholds. The fitness scores represent the log2-fold-change (LFC) between the start and endpoint gRNA abundance. LFC and quantitative genetic interaction (qGI) scores were generated as described in Aregger et al. 2020.

#### Within-vs-Between context replicate correlation (WBC)

To define within-vs-between context replicate correlations, genome-wide CRISPR-Cas9 screens completed in isogenic HAP1 cells were divided into those harboring a FASN loss-of-function mutation (n = 3) and those harboring a mutation different from FASN (n = 5), namely LDLR, SREBF1, SREBF2, ACACA and C12orf49/LUR1. At each data processing step, all pairwise Pearson correlation coefficients (PCC) within the FASN KO context and between FASN KO and each of the 5 remaining KO contexts were computed. From these comparisons, the mean 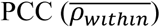 and the WBC score were computed as follows:

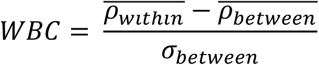

Where:

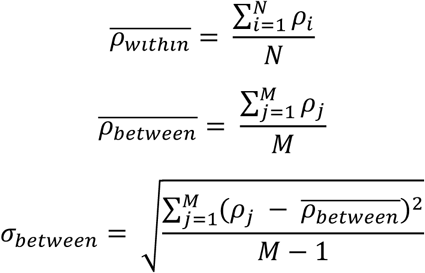

Here, N refers to the 3 possible pairwise comparisons between FASN replicated screens (within-FASN KO), and M refers to the 15 possible pairwise combinations between FASN replicated screens and LDLR, SREBF1, SREBF2, ACACA and C12orf49/LUR1 screens.

#### Cancer Dependency Map (DepMap) replicate screen comparison

LFC gRNA data (20Q3 release) was downloaded from the DepMap website [https://depmap.org/portal/]. gRNA values were mapped and mean-summarized per gene to obtain gene-level LFC data (shown in Figure 1e). Only cell lines with at least two replicates were considered in this analysis. To generate dLFC values, all screens were initially adjusted by quantile-normalizing gene-level LFC data using the R function normalizeQuantiles. Next, the scores for each gene across all screens were median centered at 0 so that if a gene was more essential in a cell line compared to its median fitness score, it had a negative score. The reproducibility of the cell line-specific signal was computed both on LFC and dLFC-level using the within-cell line PCC and the WBC comparing within-cell line PCCs to between-cell line PCCs.

#### Bin-wise replicate correlation analysis of LFC and qGI scores

To test how CRISPR screening fitness scores (LFC) affect the reproducibility of context-specific (qGI) scores, each gene was assigned to one of five bins with the intent to keep the range of qGI scores constant in each bin and having five incrementally increasing ranges of LFC scores in those bins. This was done by moving a window along the vector of qGI scores (representing the mean qGI score of both replicated screens). In each window, all genes had similar qGI but potentially different LFC scores. Genes with the most extreme LFC values were assigned to the first bin, genes with the next most extreme LFC values to the second bin and so on, thereby generating five bins with similar qGI ranges but different LFC ranges.

